# Peripheral anatomy of the dolphin ear and associated nervous structures: insights from macroscopic dissection, DICE-µCT, histology, and confocal microscopy

**DOI:** 10.64898/2026.05.15.725593

**Authors:** Steffen De Vreese, Jean-Marie Graïc, Sandro Mazzariol, Stefan Huggenberger, Marco Fogli, Federico Luzzati, Cristiano Corona, Alessandra Favole, Marc Cerdà-Domènech, Jaime Frigola, Michel André

## Abstract

The peripheral auditory system of dolphins comprises specialised bony, fatty, vascular, and neural structures adapted for underwater hearing and diving physiology. These include the external ear canal, acoustic fat bodies, sinuses, and associated neurovascular networks, which together support sound conduction, protection, and possibly sensory functions. Despite advances in gross anatomical description, the detailed integration of these tissues, particularly the innervation, neurovascular organisation, and their functional implications, remains poorly understood. Previous studies have described the presence of sensory nerve formations and vascular plexuses, but their arrangement, connectivity, and relation to each other are unresolved. Here, we combine macroscopic dissection, DICE-µCT, histology, and high-resolution confocal microscopy to characterise several neurovascular and sensory components of the dolphin peripheral auditory system in several delphinid species. Macroscopic dissection and DICE-µCT revealed the traditional acoustic fat body distribution with detailed morphology of the posterolateral extension that is not well-known. The cranial nerve distribution, and specifically the mandibular nerve branching patterns, are described in detail. Confocal microscopy uncovered a stratified neurovascular plexus around the external ear canal with a complex sensory system comprising lamellar corpuscles, Merkel cell–neurite complexes, and intraepithelial nerve fibres. Notably, the lamellar corpuscles formed a continuous, three-dimensional neural network with frequent merging and splitting of axonal bundles, shared perineuria, and vascular integration, features not observed in previous studies. Our findings demonstrate that the dolphin external ear canal and surrounding structures form a sophisticated, multimodal somatosensory organ, integrating structural, vascular, and neural specialisations likely adapted for proprioceptive mechanosensation in the aquatic environment. This study provides insights into the integration of the various components of the peripheral hearing apparatus. Future studies integrating anatomical, electrophysiological, and biomechanical approaches are needed to fully elucidate these adaptations.

## Introduction

Odontocetes (toothed whales) possess highly specialised anatomical structures in the peripheral auditory system that enable sound to be received and conducted to the middle and inner ear, collectively forming what has been termed the internal acoustic pinna (IAP) [1]. These include the mandibular bones, the intramandibular and extramandibular acoustic fat bodies, and the air-filled accessory sinus system, which together support acoustic conduction, signal modulation, and potentially other regulatory or sensory functions [2,3]. In dolphins, the original mammalian pathway for sound reception via the external ear canal is considered vestigial or functionally lost [4], while recent findings suggest that the canal may have been repurposed for alternative somatosensory roles, possibly related to diving physiology and the maintenance of auditory sensitivity under variable pressure regimes [5]. In its place, the mandibular fats, surrounding bone, and the ventral accessory sinuses form an integrated system adapted for underwater sound reception [6,7,2]. Several complementary acoustic pathways have been proposed, including transmission via the pan bone and mandibular fats [7], the gular route [8,9], lateral sound inputs [10], conduction via the dentition [11], and possible bone conduction mechanisms [12,13]. Not all of these pathways are proven and, in any case, are not mutually exclusive, rather, they appear to be frequency-dependent and spatially distributed, enabling directional hearing and spectral discrimination [2,14,15]. The structural and functional complexity of this system may also explain its vulnerability to pathological alterations observed in some stranded individuals, particularly in the context of acoustic trauma or barotrauma related to anthropogenic noise exposure or diving-related stress [16].

Despite increasing anatomical resolution, little is known about the innervation and neurophysiological integration of these peripheral tissues. The general cranial nerve architecture in cetaceans, involving cranial nerves V, VII, VIII, IX, X, and XI, retains the basic mammalian organisation but exhibits significant adaptations [17,18]. The acoustic fat bodies and the accessory sinus complex with its associated fibrovascular plexuses, including venous plexus inside the fat bodies, the sinus walls and the tympanic corpus spongiosum, are likely crucial for pressure buffering, gas exchange, and possibly thermal adaptation of acoustic properties during diving [6,3]. However, the precise anatomical integration of these structures with neural pathways and their relationship to auditory function is not yet understood [3].

Here, we investigate the macroscopic and microstructural anatomy of several components of the dolphin peripheral auditory system using a multimodal approach: macroscopic dissection, DICE-µCT imaging, histology, and confocal microscopy. We aim to clarify parts of the structural organisation, neural pathways, and potential physiological roles of soft tissues associated with the correct functioning of the peripheral auditory apparatus.

## Materials and Methods

All tissue samples were recovered postmortem from wild stranded animals and were preserved by fixation. Macroscopic dissection studies of two bottlenose dolphin (*Tursiops truncatus,* Montagu, 1821) and Pantropical spotted dolphin foetus (*Stenella attenuata*, Gray, 1846) were performed, the former during routine necropsies at BCA, UniPd, and the latter performed on a museum specimen at the Department II of Anatomy (Neuroanatomy), University of Cologne (50924 Cologne, Germany). Other macroscopic and microscopic samples were obtained from bottlenose and striped dolphins found stranded along the Italian and Spanish coastline and a full post-mortem examination following an internationally standardised procedure [19] was performed by trained personnel aiming at determining the cause of death and health status (Details can be found in [5]). Standard macroscopic dissection was carried out to expose ear-related tissues, muscles, sinuses, and neural components. Detailed macroscopic studies of the cranial nerve distribution were performed, with special attention to the innervation of the peripheral auditory apparatus and associated structures. For this, a single Pantropical spotted dolphin foetus (*S. attenuata*) that had been fixed in formalin for more than 40 years, was dissected macroscopically to map the cranial nerves associated with the region of the outer ear canal and tympanic membrane.

For histological analysis, selected samples from various animals were fixed in 10% neutral-buffered formalin, paraffin-embedded, sectioned, and standard HE staining was applied to 50 consecutive 5-µm transverse sections spanning a 400-µm length of the external ear canal wall in a striped dolphin specimen. Digitised images were cropped centred on the canal and aligned (rotation and translation) using the TrakEM2 plugin for ImageJ [20], in combination with the Virtual Stack Registration plugin [21] where necessary, followed by manual correction of the alignment. Nervous structures were labelled manually without distinction between sensory nerve endings and nerve fibres, as HE staining did not allow unambiguous differentiation between lamellar corpuscles and nerve profiles. Labelled coordinates were exported as a CSV file and imported into Blender (v2.79–2.82; Blender Foundation) using a custom Python script for 3D volume rendering and visualisation.

For confocal microscopy, formalin-fixed, paraffin-embedded samples from the out ear canal of a dolphin were sectioned at 5 µm using a microtome. A total of 60 serial sections were mounted on positively charged glass slides and dried overnight at 37 °C. For immunofluorescence analyses, a serial staining approach was applied: three consecutive sections were subjected to double labelling with NF and PGP9.5, while one section every three serial sections was stained with NF and S100. For Immunofluorescence staining, sections were deparaffinized using Ottix Plus and Ottix Shaper (Diapath, Bergamo, Italy), rinsed in running tap water, and subjected to antigen retrieval in citrate buffer (pH 6.1) at 95 °C for 20 min. Sections were then incubated in distilled water for 5 min, followed by washes in PBST (phosphate-buffered saline containing 0.05% Tween 20). To block non-specific binding, sections were incubated for 1 h at room temperature in 5% normal donkey serum with 0.3% Triton X-100 (Merck, Darmstadt, Germany), and then incubated for 1 h at room temperature in a humid chamber with the following primary antibodies diluted in PBST containing 0.3% Triton X-100: polyclonal anti-bovine S-100 (Dako, Z0311; RRID:AB_10013383; 1:400), monoclonal anti-human neurofilament M (NF; Dako, M0762; RRID:AB_2314899; 1:100), and polyclonal anti-bovine PGP9.5 (Dako, Z5116; RRID:AB_2622233; 1:100). After washing, sections were incubated for 1 h at room temperature in the dark with fluorophore-conjugated secondary antibodies: Alexa Fluor 488 donkey anti-mouse IgG (Thermo Fisher Scientific; RRID:AB_2535792; 1:1000) and Alexa Fluor 647 donkey anti-rabbit IgG (Thermo Fisher Scientific; RRID:AB_2535813; 1:800). Slides were washed in PBST and mounted with ProLong Gold Antifade Mountant with DAPI (Thermo Fisher Scientific; RRID:SCR_015961). Negative controls were performed by omitting the primary antibodies. Antibodies were previously validated in [5].

For DICE-µCT analysis, samples from the striped dolphin specimens originated from the Spanish stranding network, and obtained during routine necropsy (n=2, *Stenella coeruleoalba*, Meyen, 1833). After removal of the tongue, ventral part of the larynx and the tongue bones, a block of tissue centred around the TPC, including soft tissues and bones of the skull and mandible, was dissected with knives and saw, keeping the anatomy of the region as intact as possible. Samples were fixed in 10% neutral-buffered formaldehyde before being iodine-stained (I_2_KI 2% concentration with three liquid changes during 34 months), and then washed with distilled water and scanned using high-resolution X-ray micro-Computed Tomography (μCT, MultiTom Core from XRE – X–Ray Engineering, nowadays TESCAN) at the CORELAB Laboratory of the University of Barcelona. The scan of samples N101-21 and N52-21-02 was performed at 160 kV, 58 W, with no filter, exposure time of 160 ms, 2 averages, and 1001 projections, resulting in a voxel resolution (volumetric pixel or cubic unit representing a 3D object) of 58 μm. While the scan of sample N52-21-01 was performed at 160 kV, 55 W, with a 0.5 mm Cu-filter, exposure time of 250 ms, 2 averages, and 1001 projections, resulting in a voxel resolution (volumetric pixel or cubic unit representing a 3D object) of 55 μm. Acquisition and reconstruction of 3D raw data were carried out with Acquila and RECON software packages from XRE). Digital tissue segmentation was conducted using 3D Slicer (v5.9.0, slicer.org)[22] using manual and semi-automated tools, segmentations exported as STL files, postprocessed in Meshlab [23], and imported into Blender (v4.4.1)[24] for visualisation.

## Results

The following results describe the peripheral anatomy of the dolphin auditory region, integrating macroscopic dissection, DICE-µCT, histology, and confocal microscopy to reveal the complex interplay of mechanoacoustic, vascular, and neural structures. The findings are organised into functional anatomical systems, acoustic pathways, pressure-modulating structures, and the ear canal and middle ear, to highlight their distinct morphologies and interactions.

### Region of the ear canal and middle ear

In the macro-and microscopic analyses, the external ear canal exhibited the distinctive downward spiral and followed a consistent topographical pathway: running horizontally through the blubber, then turning at a straight angle ventrally, then continuing as a mediorostroventrally spiralling canal accompanied by cartilage, before arriving more or less straight horizontally at the tympanic (Fig 1). The ear canal passes ventral to the bilobed middle sinus and anterior to the posterior sinus. In the most medial couple of centimetres, the canal’s wall consisted of soft tissue of comparable radiodensity to the acoustic fat bodies. In one specimen, the right ear canal contained a discrete air bubble at its medial end, not associated with any observable artefactual connection to the external environment, while the left ear canal of that same animal showed air in its medial third, likely attributable to a dissection artefact. The second specimen showed no in-situ air in either canal, yet the canal remained clearly visible due to the lower tissue density relative to the surrounding connective tissue. Adjacent to the canal, a thin cartilaginous structure, in DICE-µCT only distinguishable in its medial half by its shape and known histological location, ran parallel to the canal from the ventral spiralling down to the entrance to the middle ear (Fig 2).

**Figure 1:**
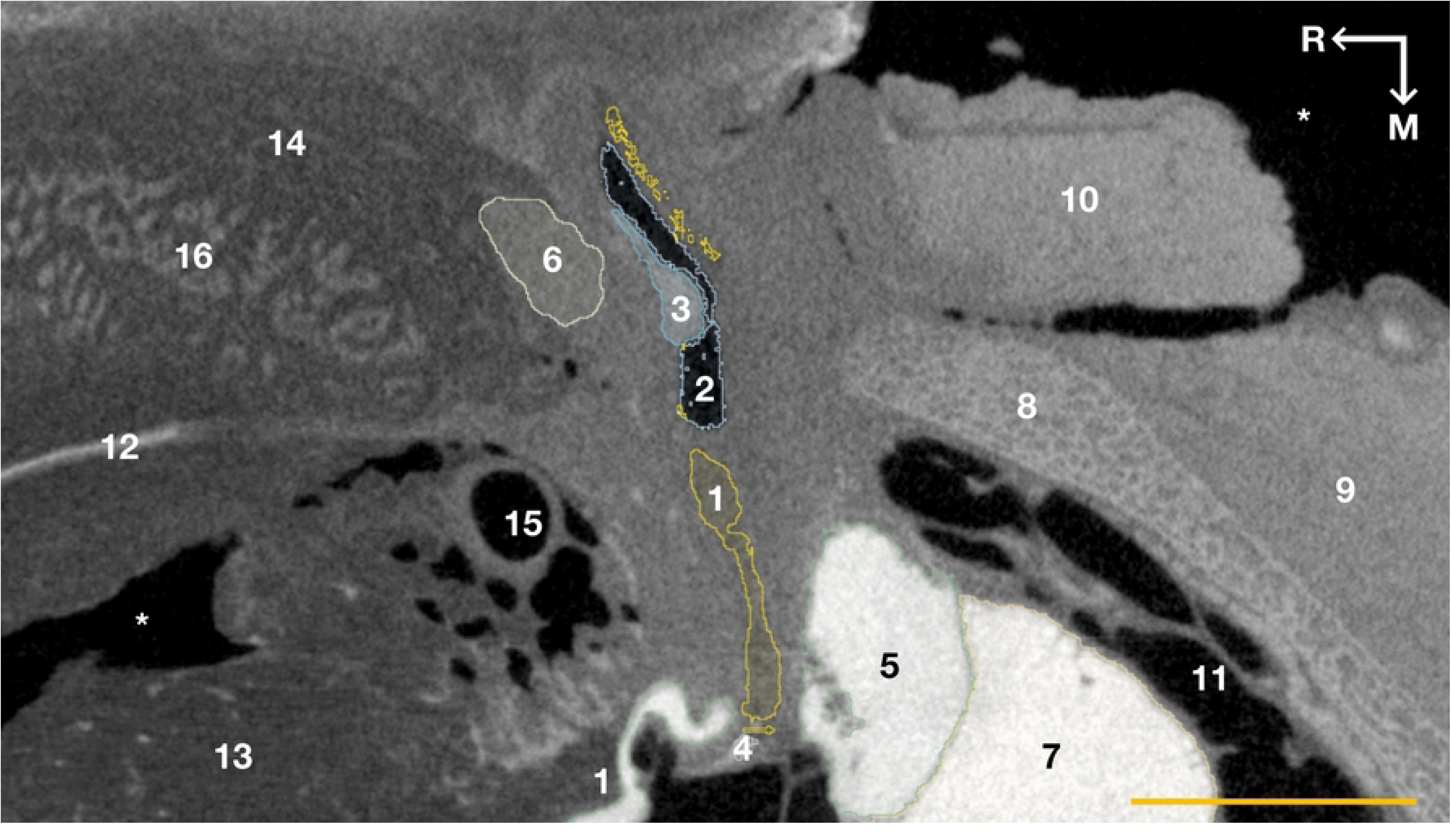
DICE-µCT sagittal section at the level of the left ear structures. The ear canal (1) with its (artefactual) lumen (2) and the nervous tissue ridge (3) that is present in the medial third of the ear canal. The ear canal ends in a content-filled bulbous inflation sustained in the cup of the tympanic membrane (4), which continues as the tympanic ligament and which itself shows a small soft tissue connection to the posterior process of the tympanic bone (5). The facial nerve (6). The tympanic and periotic (7) bones. The skull exoccipital bone (8), and the scalenus (9) and sternocephalic (10) muscle. The peribullar sinus (11) with air, ligaments and vascular structures. The mandibular bone (12), the IMFB (13) and EMFB (14). The vascular plexus of the IMFB with the large maxillary artery (15). The masseter muscle (16) intertwined with the EMFB. Asterisks: artefactual space. 10 mm scale bar. Arrows indicate rostral (R) and medial (M) directions.

**Figure 2:**
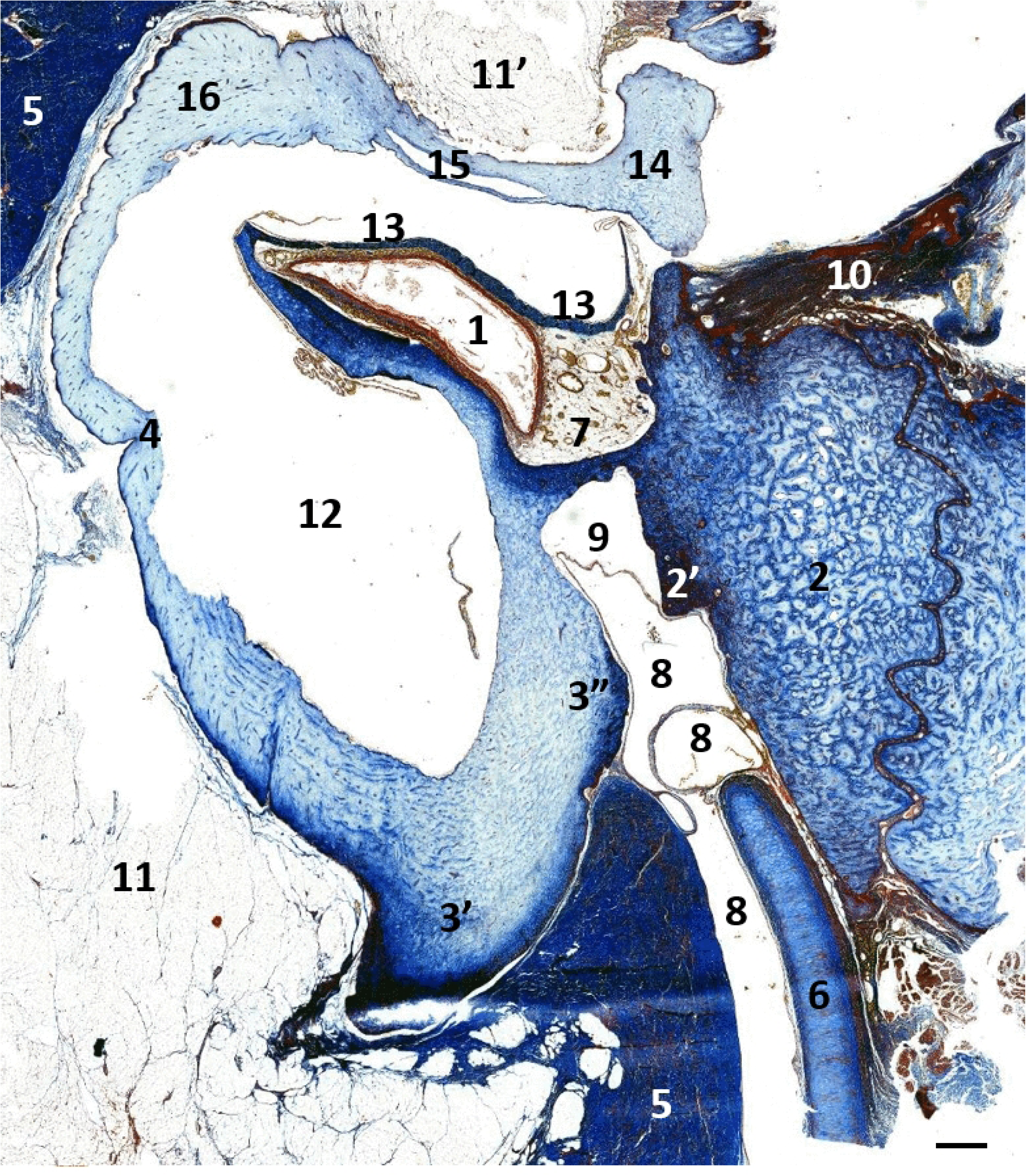
Masson’s trichrome staining of the left ear canal entering the TP-complex at the level of the tympanic notch in striped dolphin. 1. Ear canal; 2. Posterior process of the tympanic (Mead and Fordyce, 2009) or processus petrosus (Nummela et al., 1999); 2’ posterior tympanic spine; 3. Sigmoid process of the tympanic; 3’ dorsal apex; 3” epitympanic border; 4. Sulcus for chorda tympani; 5. Connective tissue capsule surrounding the acoustic fat; 6. Ear canal cartilage; 7. Vascular network in a fibrine mesh; 8. Vascular sinuses accompanying the ear canal on its course through the tympanic aperture; 9. Artefactual space; 10. Ligament attaching to 2, for suspension of the TP complex in the peribullar sinus; 11. Acoustic fat contacting the sigmoid process and the malleus (11’); 12. Tympanic cavity projecting into the sigmoid process; 13. Tympanic membrane; 14. Malleus; 15. ‘blue dot in Cranford (2010, Fig 10); 16. Accessory ossicle. Scale bar 1mm.

Several auricular muscles were identified in association with the external ear canal. A thin muscle layer corresponding to the occipito-auricularis muscle was observed just beneath the blubber, inserting caudally into the connective tissue surrounding the canal along the vertical part of its course. This muscle was continuous with the occipito-auricularis superficialis, accompanying the ear canal laterally into the blubber. The zygomatico-auricular muscle was also noted to insert into the ear canal tissue at the level of the ventral curvature, but on its rostral surface, and its innervation by several branches of the facial nerve was confirmed through DICE-µCT (Fig 3). Muscle nomenclature follows the terminology used by Purves [25].

**Figure 3.**
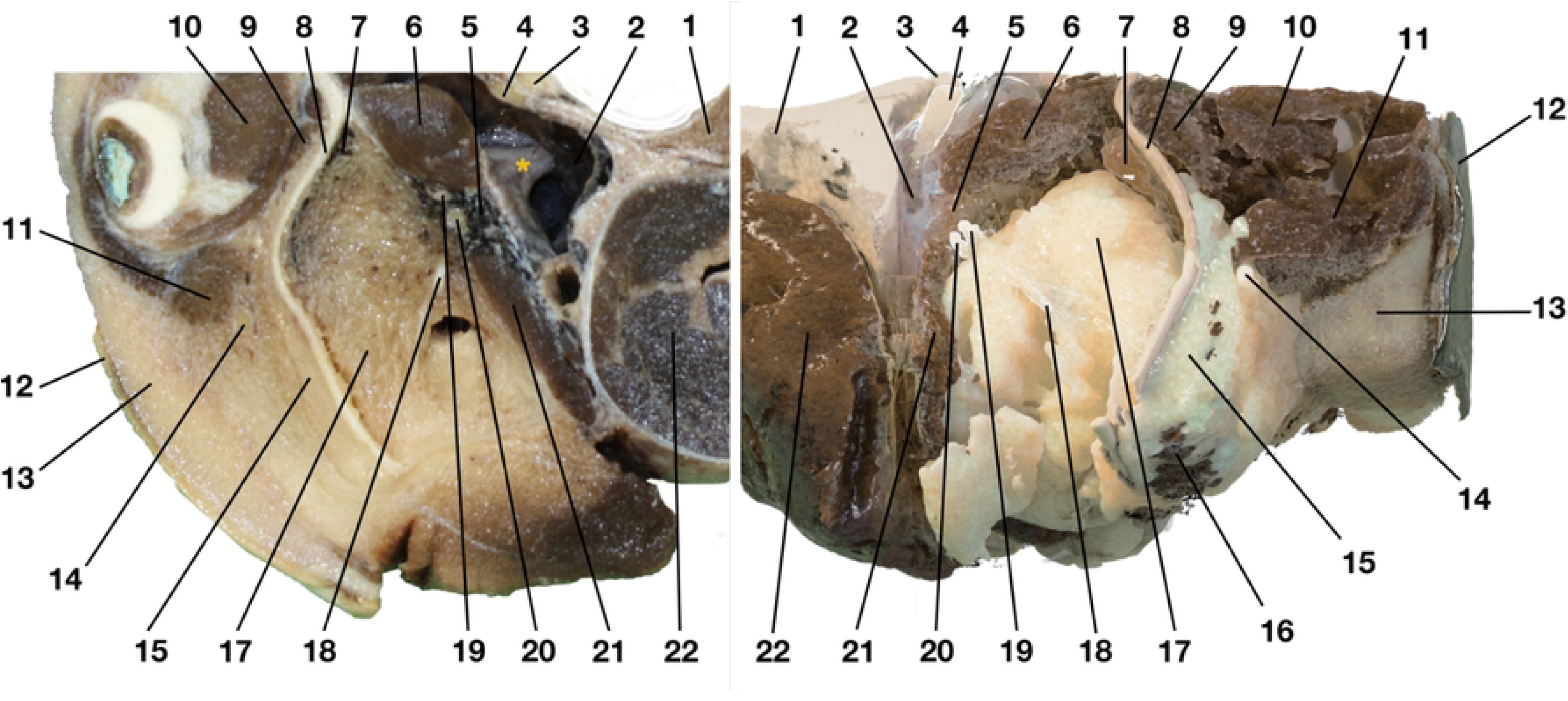
Comparative view of a macroscopic rostral photograph of a transverse section through the head of a striped dolphin on the left, compared with a 3D reconstruction from DICE-µCT of the right side (same species, different animal). Mandibular nerve (yellow asterisk) exiting from the cranium into the pterygoid sinus (2), branching into the monogastric (18) and mylohyoid and inferior alveolar nerves (19-20) on their laterorostroventral course, entering the lateral wall of the pterygoid sinus in between the lateral and medial pterygoid muscles (6, 21) and surrounded by part of the pterygoid vascular plexus (5). 1. Skull, 3. Optic nerve, 4. Opthalmic and maxillary nerve, 7. Vascular plexus on inside of the mandibular ramus, 8. Mandible, 9. Temporal muscle, 10. Eye muscles, 11. Zygomatico-auricular muscle, 12. Skin, 13. Blubber, 14. Facial nerve, 15. EMFB, 16. Masseter muscle embedded in the EMFB, 17. IMFB, 21. Medial pterygoid muscle, 22. Pharynx and associated musculature

Regarding innervation of the ear canal region, the mandibular nerve (V₃) was traced from its exit from the foramen ovale at the base of the skull to its auriculotemporal branch, which arose from the mandibular ganglion near the branching point of the monogastric nerve. The auriculotemporal nerve coursed caudolaterally through the IMFB toward the ear canal and bifurcated into two principal branches along its rostral wall (Fig 3). These gave rise to several smaller fibres, which became too fine to trace macroscopically and did not show significant contrast differences in DICE-µCT. On passing the caudal border of the mandible, we noted indications of a fine nerve branch connection to the facial nerve. The dense fibrous tissue between V₃ and the medial end of the external ear canal was notably resistant to macroscopic dissection, as also shown in histology, while DICE-µCT scans confirmed these observations. The auriculotemporal nerve and its branches were visualised as paired neurovascular bundles enveloped in connective tissue, in close interaction between IMFB, mandible, temporomandibular joint, and the mandibular vascular plexus (Figs 4 and 5).

**Figure 4.**
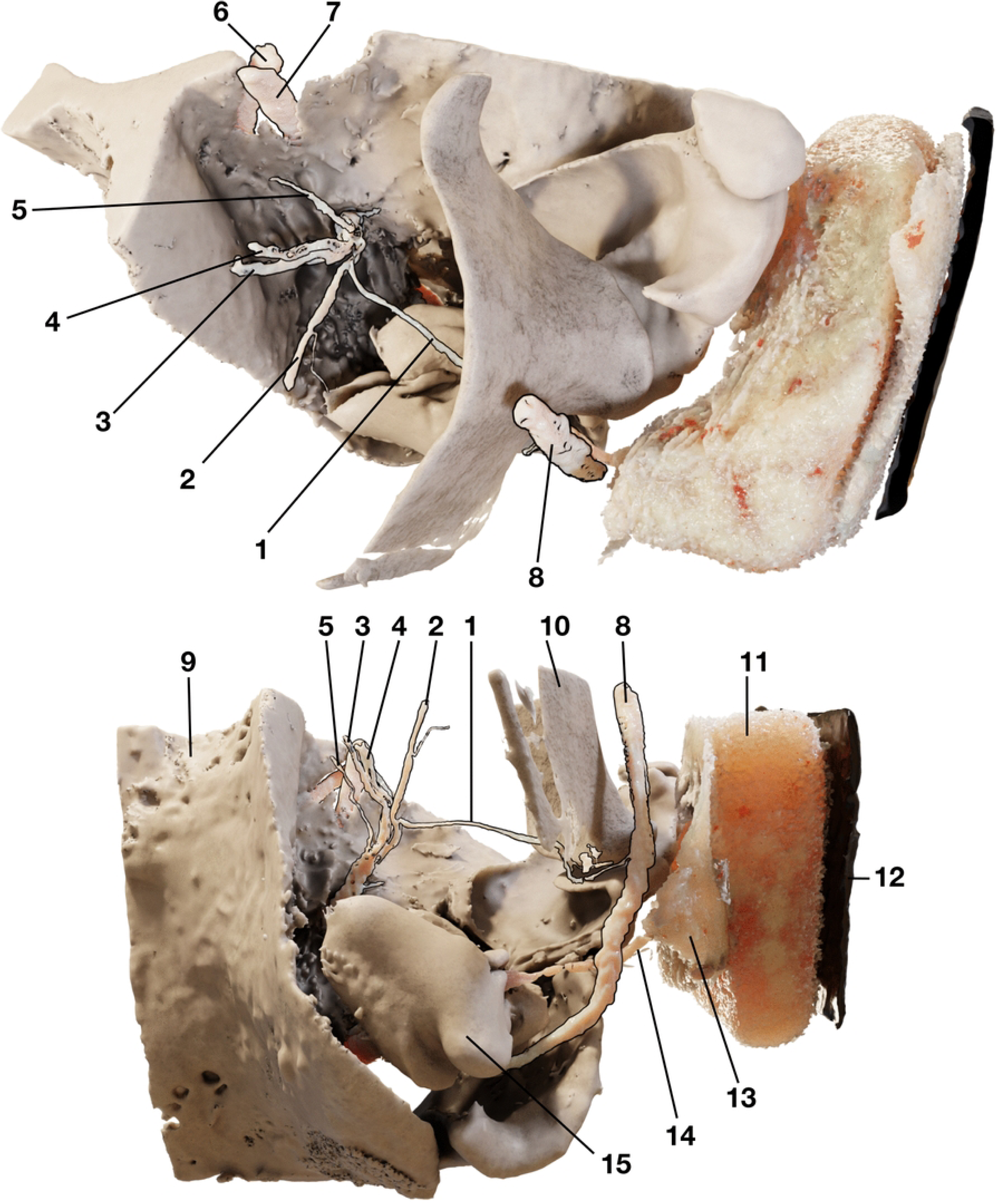
Comparative view of the left mandibular and facial nerve topology in 3D reconstruction of the ear region. Top: rostrolateral view, Bottom: ventral view. 1. auriculotemporal nerve, 2. monogastric nerve with a small ventral branch. 3. mylohyoid nerve, 4. inferior alveolar nerve, 5. pterygoid nerves, 6. optic nerve, 7. ophthalmic and maxillary nerve, 8. temporal nerve, 9. facial nerve, 10. skull, 11. mandible, 12. blubber, 13. skin, 14. posterolateral extension of the EMFB, 15. external ear canal, 16. tympanic bone.

**Figure 5.**
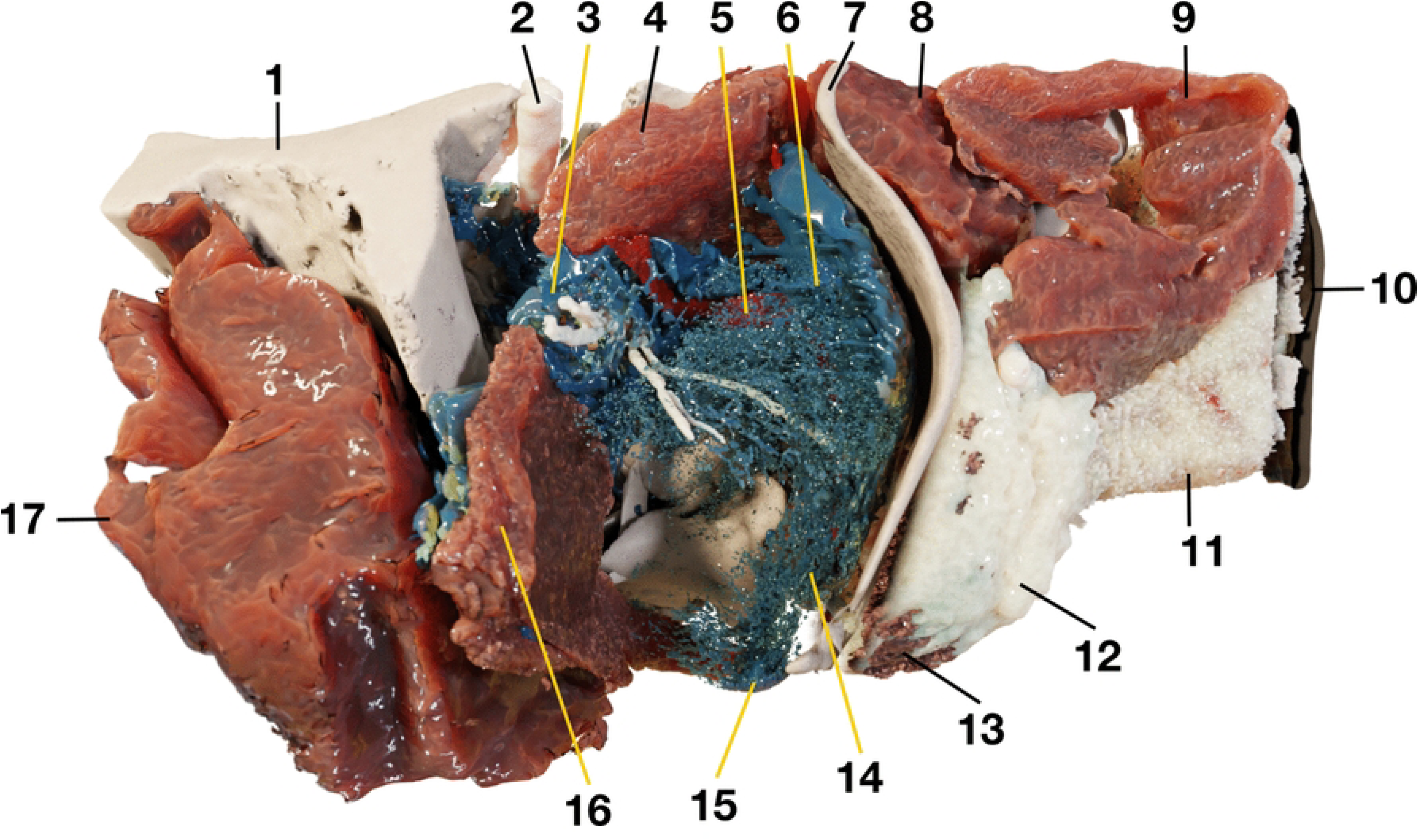
Rostral view of a 3D reconstruction of the left ear region. Left: the IMFB and pterygoid sinus plexuses (blue) extending from the tympanic bone along medial wall of the mandible and, accompanying the maxillary vein (red), running dorso-caudally around the IMFB between lateral and medial pterygoid muscle and through which the inferior alveolar nerve and the monogastric nerve pass. Vascular tissue inside the IMFB is mainly present in the laterodorsal half of the IMFB, being very sparse medioventral region, forming a continuous cylindrical network that extends from rostral to caudal where the IMFB contacts the tympanic bone. Also note the vascular plexus on the medial wall of the pterygoid sinus, visible medial to the medial pterygoid muscle. 1. skull; 2. optic and ophthalmic nerve; 3. pterygoid vascular plexus; 4. lateral pterygoid muscle; 5. maxillary vein; 6. continuation of the pterygoid vascular plexus in the pterygoid sinus; 7. mandible; 8. temporal muscle; 9. zygomatic muscle with part of the posterior muscles of the eye; 10. skin; 11. blubber; 12. EMFB; 13. masseter muscle; 14. IMFB vascular plexus; 15. tympanic bone; 16. medial pterygoid muscle; 17. pharyngeal constrictor muscle.

Further medial macroscopic dissection revealed additional cranial nerve components within the middle ear. From the mandibular ganglion branched a nerve observed to enter the middle ear cavity through the anterior incisure of the periotic bone, running between the tympanic and periotic towards the malleus. Two caudally directed branches, one dorsal and one ventral, arose from the mandibular ganglion and appeared to contribute to the middle ear innervation, with the dorsal branch being more prominent. The vagus nerve (X) was identified macroscopically, but no branches were observed. The recurrent laryngeal nerve, running alongside the hypoglossal nerve (XII), was also identified. The chorda tympani branch of the facial nerve was not observed with any of the applied methods.

In confocal microscopy of a 300 micron section of the external ear canal of striped dolphin, the sensory and neurovascular architecture revealed a highly complex and interconnected organisation, integrating lamellar corpuscles, nerve fibres, Merkel cell–neurite complexes (MCNs), and vascular components into a circummeatal network with multiple levels. 2D and 3D analysis demonstrated an extensive intramural nervous plexus surrounding the canal’s epìthelium, in which nearly all identified nervous structures formed a continuous web. Few apparently separate structures, not visibly connect to the main plexus in the observed planes, may have been incompletely captured due to the limited size of the sample, and could possibly be connected to the nervous network outside the sample range.

The ear canal exhibited two distinct neurovascular plexuses: a deep neurovascular plexus (DNP) located ∼500–700 µm from the canal centre and a superficial neurovascular plexus (SNP) situated ∼150 µm beneath the epithelium (Fig 6). These plexuses ran parallel to the canal, forming a circummeatal sensory web with embedded vascular elements. High-density innervation was noted around arteries and arterioles, with NF-immunoreactive axons accompanying arterial walls and PGP9.5 showing patchy staining of vessel walls. In at least two instances, blood vessels were enclosed within the perineurium of nerve bundles.

**Figure 6.**
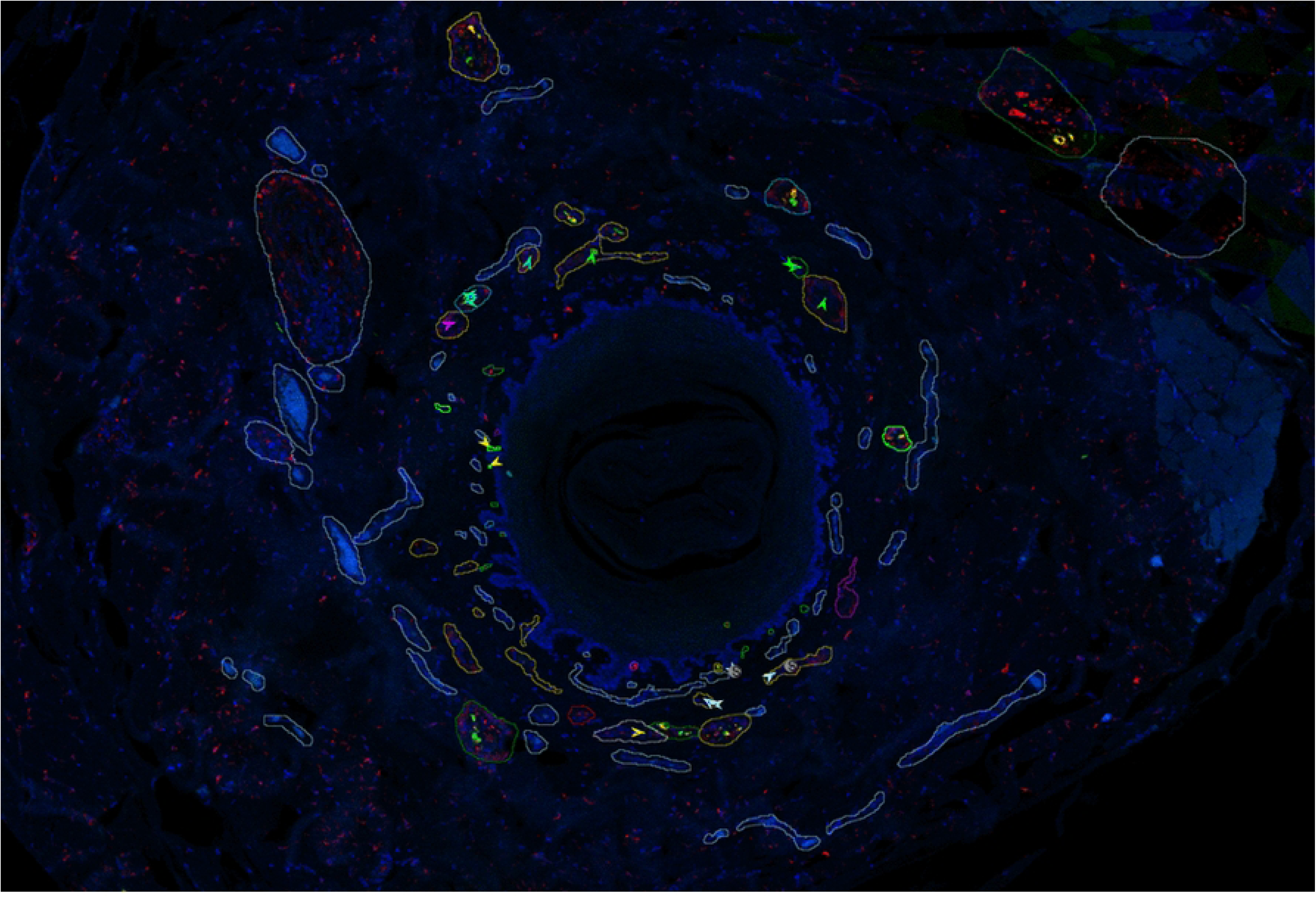
Confocal microscopy transverse section (anti-DAP1 = blue, anti-PGP9.5 = red, anti-NF = green (and also yellow where overlap with anti-PGP9.5) through the ear canal of a striped dolphin at about 2 cm beneath the skin. Deep and superficial neurovascular plexus, and connection between the two. On the top right are visible a large artery (1) and nerve (2), part of the deep neurovascular plexus, providing innervated and vascularized connections (3) to the superficial neurovascular plexus in which there are smaller vascular structures (white outline lables) and neural structures (various coloured outline labels) including lamellar corpuscles and extensions into MCN’s.

Lamellar corpuscles were consistently located in the subepithelial layer, longitudinally oriented, and originating from small nerves with few axons. In some cases, the nerve appeared to transgress entirely into one or multiple corpuscles, while in others, corpuscles branched off from nerves that continued distally, although still embedded within the same perineurium, forming a local ‘receptor-nerve-complex’ (Fig 7). Corpuscles showed frequent bifurcations and convergences as they coursed toward the periphery. Corpuscles rapidly increased in diameter after branching from a nerve, with lengths ranging 100–500 µm, generally elongated, with lamellar structure covering most of the corpuscle. In several examples, corpuscles curved up to 180°, changed direction, or continued and merged back into the neural plexus, forming a continuous network rather than isolated sensory endings.

**Figure 7.**
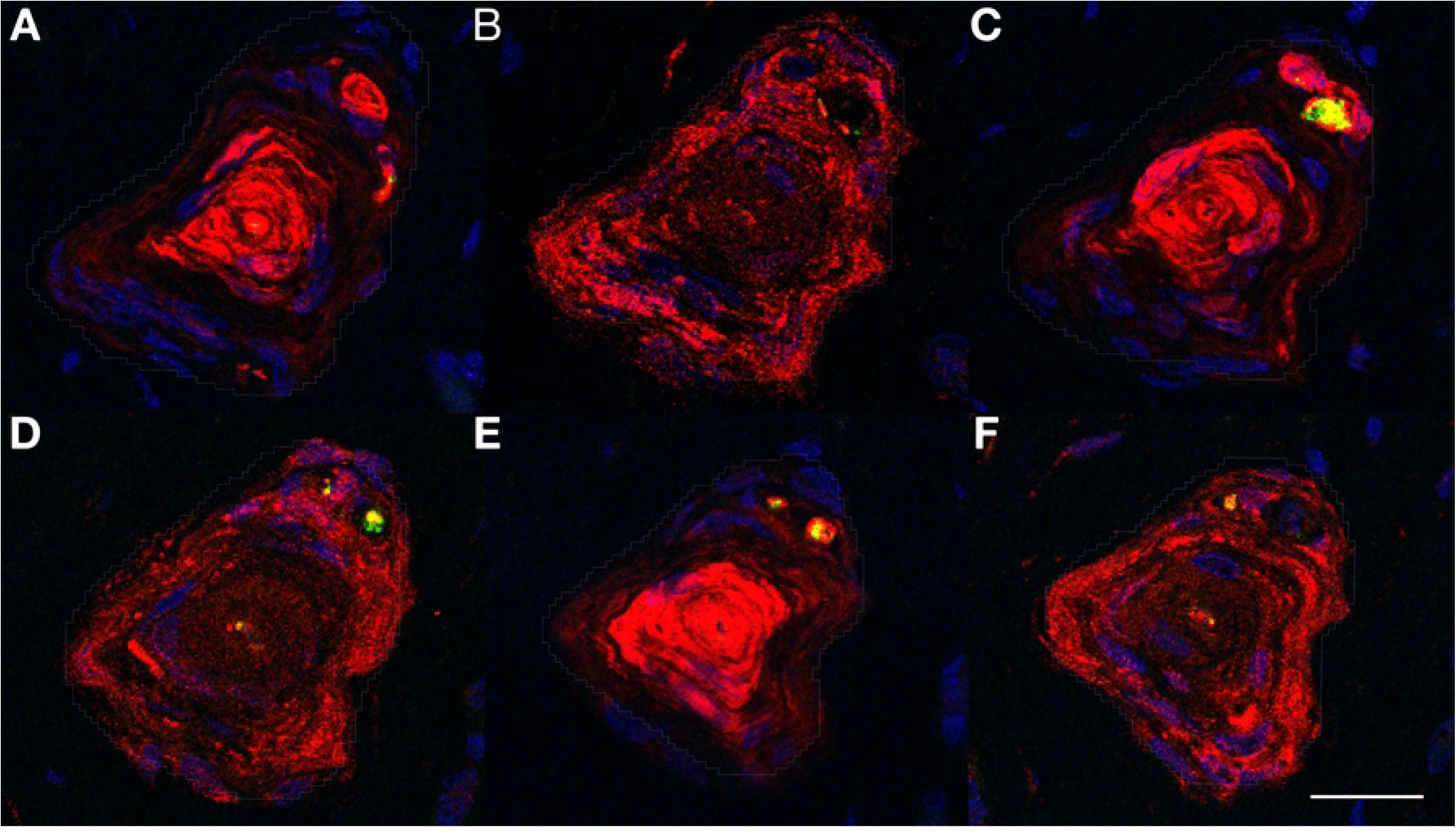
Five consecutive images alternatively stained with a combination of NF, Dap1, and S100 (A,C,E), and NF, DAP1, and PGP9.5 (B,D,F) showing consective sections through a lamellar corpuscle with adjacent nerves in a shared perineurium. Immunoreactivity to anti-NF shows the central axon in the corpuscle, and the adjacent nerve fibres. Anti-S100 stains the inner core lamellae of the corpuscle, and the myelin sheath aroud the nerve axons. Anti-PGP9.5 immunoreactivity is visible in the outer lamellae of the corpuscle, less intense also in the inner lamellae, and the perineurium of adjacent nerves. Scale bar 20 µm.

Beyond lamellar corpuscles, we identified MCNs in the basal lamina of epidermal ridges at the dermal–epidermal junction. These comprised a subepithelial afferent neurite (NF+, PGP9.5+) in contact with a PGP9.5+, S100– Merkel cell, consistent with MCNs in terrestrial mammals. Intraepithelial nerve fibres (IENFs) were observed within the stratum spinosum, with some structures positive for NF and others for PGP9.5. We also detected previously undescribed intraepithelial PGP9.5+ structures lacking NF and S100 immunoreactivity, suggesting an unidentified sensory or glial component. One unique structure was positive for both S100 and NF, potentially consistent with a Langerhans cell. Overall, the confocal data revealed a highly interconnected neural and vascular network, integrating classic mechanosensory elements and novel structures into a complex circummeatal sensory system.

The 3D reconstruction of the intramural nervous plexus over a 400-µm canal segment revealed a highly interconnected neural network in which almost all identified nervous structures formed part of the same continuous sensory complex (Figs 8-9). The few structures not connected to the principal network within the reconstructed volume are not necessarily isolated, as connections may exist in sections beyond the sampled length. The reconstruction demonstrated that the plexus extends throughout the full circumference of the canal wall and that the lamellar corpuscles and nerve profiles are integrated into a single, spatially continuous system. Although HE staining did not allow for unambiguous distinction between individual corpuscles and nerve fibres, similar to confocal microscopy showing transitory morphologies, while the spatial continuity and density of the labelled structures were consistent with the rich subepithelial neurovascular architecture described in confocal microscopy sections of the same region. This partial reconstruction is considered representative of the general plexus organisation in the distal segments of the ear canal, while the medially located nervous tissue ridge, as documented earlier [26], was not included for analysis.

**Figure 8.**
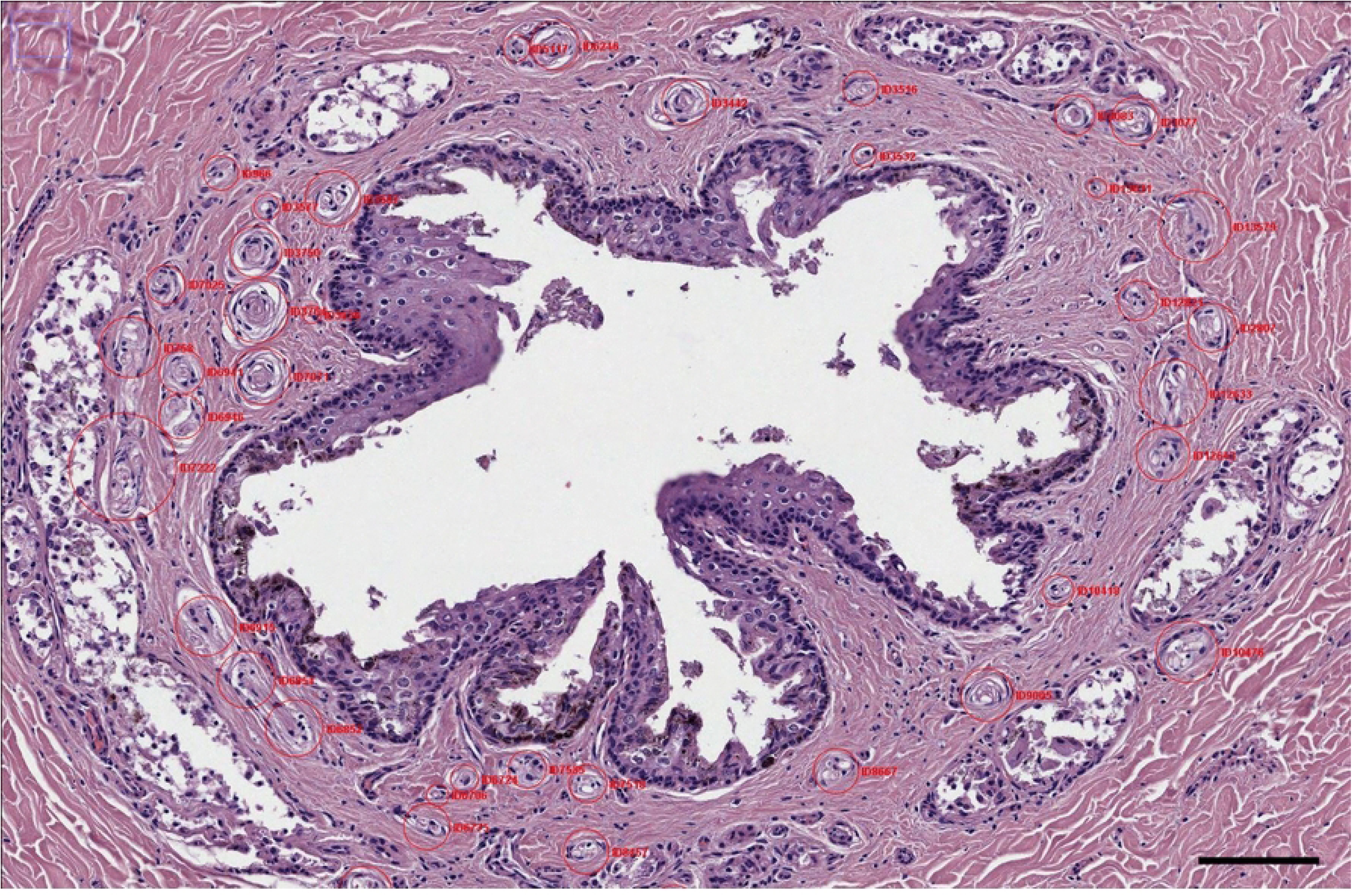
Histological image (HE staining, striped dolphin ear canal superficial third) of a single histological section. The Trackmate plugin overlay shows manually labelled nervous structures encircled in red with individual ID number. Scale bar 100 µm.

**Figure 9.**
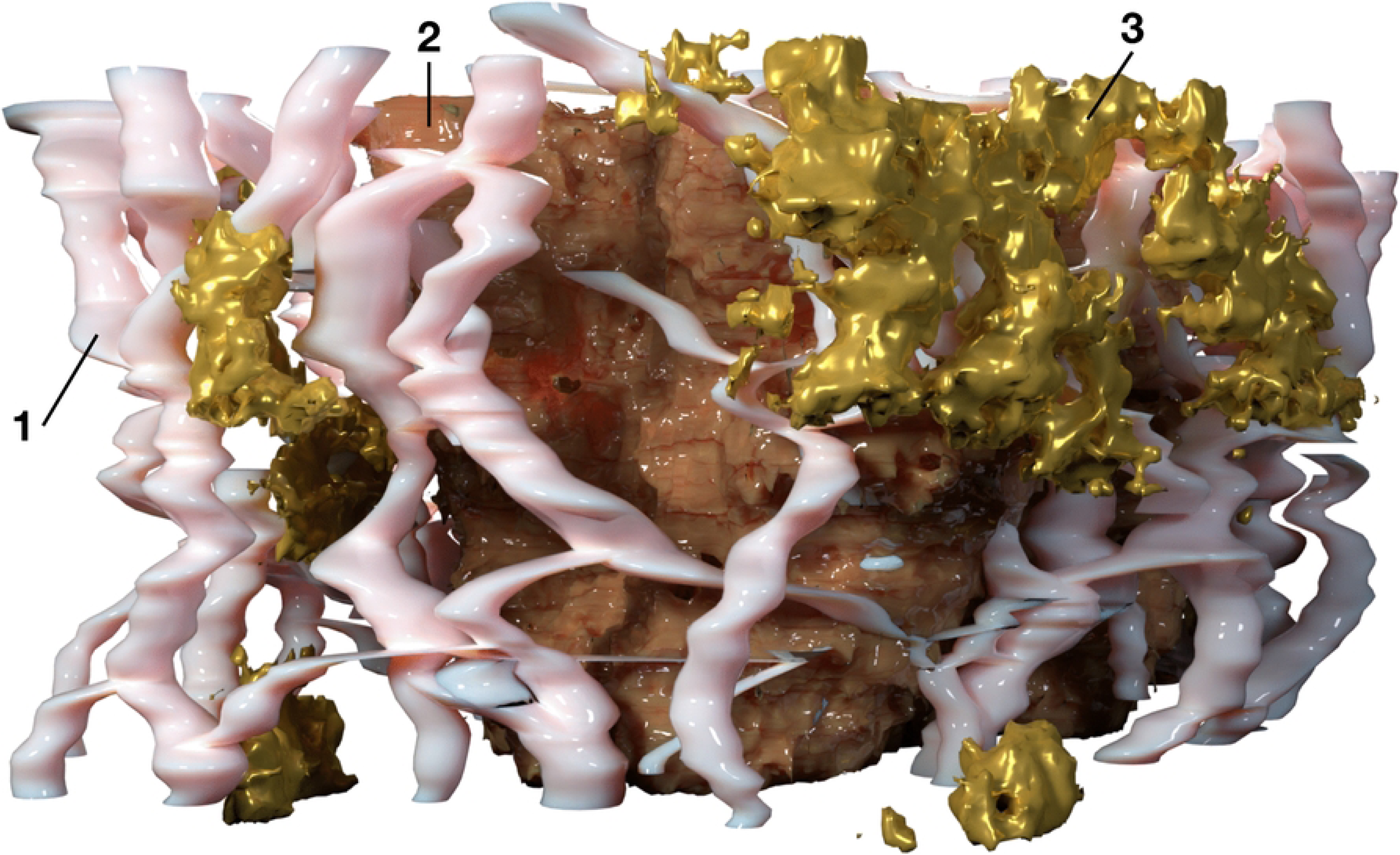
3D volume rendering based on histological sections of the external ear canal (positioned vertically, 400 µm length). The intramural nervous plexus (1) forms a continuous network around the ear canal epithelium (2) and is intricately associated with the ear canal glands (3) at this level of the ear canal.

### Mandibular fat bodies

The acoustic fat bodies presented a highly organised configuration with the intra-and extramandibular fat bodies forming a continuous structure. The IMFB occupied the mandibular canal and extended caudomedially through the mandibular foramen, merging with the extramandibular fat body (EMFB) caudal to the mandibular ramus and ventral to the temporomandibular joint. The EMFB was situated lateral to the mandible, overlaying the pan bone, and curving caudally and slightly ventrally around the caudal ramus. Together with the IMFB, the internal end of the fat body complex bifurcates, with one branch inserting into the acoustic ear funnel located between the periotic and tympanic bones, and another attaching to the outer surface of the tympanic plate. At its caudolaterodorsal aspect, the EMFB also contacted the external ear canal-associated fat and adjacent blubber, forming a posterolateral end. This distinct posterolateral extension projected laterocaudally toward the external ear canal, running parallel to the canal along its sigmoid course, positioned approximately 5–10 mm rostral to it, and with indications of flaring outward like a trumpet where it reached the blubber layer encompassing the ear canal’s emergence from the blubber, although tissue artefacts of sample preparation did not allow for a complete dissection (Figs 1 and 4). These morphological findings suggest a structural arrangement of the fat bodies that integrates the mandible, the middle ear and the external ear canal, forming a spatially continuous system that interfaces with peripheral nervous structures.

The acoustic fat bodies exhibited multiple nervous structures closely associated with their surfaces and traversing through their substance, as observed macroscopically, in DICE-µCT scans, and histologically. Detailed dissections and DICE-µCT revealed multiple ramifications of the mandibular nerve (V_3_) as it emerged from the skull and bifurcated into several branches, including the auriculotemporal, monogastricus, mylohyoideus, inferior alveolar, and smaller nerves presumed to be pterygoid branches (Fig 4). The auriculotemporal nerve branched off approximately 10 mm distal to the bony foramen, coursing rostroventrally together with the monogastric nerve for a short distance, before turning laterocaudally toward the external ear canal.

The monogastric nerve continued rostroventrally into the IMFB, where it gave off a small ventral branch. The inferior alveolar and mylohyoid nerves were observed running parallel (mylohyoid nerve presumably medially), passing through the conspicuous pterygoid vascular plexus situated between the IMFB laterally, the medial pterygoid muscle ventrally, the lateral pterygoid muscle dorsally, and the pterygoid sinus medially (Fig 5). Measurements of the main nerves in the ear region showed diameters of about 0.7 mm for the auriculotemporal, 1.0 mm for the inferior alveolar and mylohyoid, 0.5 mm for the pterygoid nerves, 1.2 mm for the monogastric, and 3.0 mm for the facial nerve. The EMFB, with the embedded masseter muscle showing intertwining of fat and muscle tissue, also contained nervous structures. While, the facial nerve traverses the EMFB, running lateral to the masseter muscle and giving off a few small branches towards the zygomaticoauricular muscle, histologically, there were also smaller nerve fibres and occasional lamellar corpuscles within the EMFB’s sparse but well-vascularized connective tissue. Similarly, in the IMFB, medium-to-large nerve fibres were observed traversing, together with indications of a few terminal sensory structures such as lamellar corpuscles in the sparse connective tissue.

Both histological and DICE-µCT analyses revealed a dense, continuous microvascular network throughout the IMFB, with a similar but less prominent vascular investment in the extramandibular fat. Small veins and arteries were observed penetrating deeply into the fat body, arranged as an interconnected, multidirectional network rather than confined to the periphery. Vascularisation was sparser in the IMFB’s central core, compared to more peripheral, and the core continued caudomedially to contact the TP complex in previously described locations. Intricately associated with the IMFB plexus, there was a localized vascular plexus, part of the pterygoid plexus, situated on the dorsocaudal margins of the IMFB, that involves the IMFB from the medial extension between pterygoid sinus and IMFB, accompanying the maxillary vein, revolving caudally around the IMFB and passing rostrally to the middle sinus at the level of the temporomandibular joint while also continuing at the caudodorsal margin between the IMFB and the mandibular bone in direction of the tympanic bone.

The masseter muscle was identified as a reduced structure originating from the zygomatic arch and inserting on the lateral surface of the mandible near the gonial angle. Muscle fibres were sparse and interspersed with adipose tissue, and the masseter was closely associated with the venous and fat body networks. The masseter nerve was not identified.

The walls of the pterygoid and peribullar sinuses contained numerous large vessels embedded in dense connective tissue, as well as several small to medium-sized nerve fibres and a ganglion within the medial wall of the pterygoid sinus near the opening of the tuba auditiva. Additional nerve fibres were noted in the ventrolateral pterygoid sinus wall at the interface with the IMFB, often in proximity to vascular structures. Sparse corpuscles were occasionally seen, particularly near vessels, but no specialised sensory nerve formations were detected. Arteries were also identified in the walls of the sinuses and within the IMFB, where they coursed alongside the venous network, although their full distribution and branching patterns could not be traced in the available sections.

### 1. Accessory sinus system and vascular plexuses

Trabeculated air spaces were observed bordering the pterygoid, sphenoid, and occipital bones, and the tympano-periotic complex. These spaces included the pterygoid sinus, posterior sinus, and middle sinus. The walls of these sinuses abutting the bones and the trabeculae connecting opposing bones consisted of intensely vascularized connective tissue, consisting of both arterial supply and extensive venous networks. In microdissection, the pterygoid venous plexus was empty of blood, making it difficult to distinguish veins from the extensions of the air spaces. The larger, localised, venous plexuses included the intramandibular venous plexus, peribullar venous plexus, pterygoid venous plexus, and the vascular network of the corpus spongiosum tympanicum. The intramandibular venous plexus was located within the IMFB, forming a dense network along the mandibular canal except for a central region, slightly ventral, that continued caudally to contact the middle ear, in which the vascular network was not obvious on DICE-µCT (see acoustic fat bodies). The peribullar venous plexus surrounded the tympano-periotic complex, while the pterygoid venous plexus was embedded within the walls of the pterygoid sinus and between the pterygoid muscles, and continued along the pterygoid bones along the region of the Eustachian tube as the peritubal venous plexus and abutting the pharyngeal musculature. The Eustachian tube itself was tracked from the nasopharynx to the peribullar cavity, opening slightly anterior to the tympanic bulla and its opening to the tympanic cavity. The EMFB was vascularized but did not present an extensive vascular plexus as in the IMFB. The corpus spongiosum tympanicum was situated within the tympanic cavity, visible as a highly vascular structure continuous with the walls of the peribullar sinus, and containing the central internal carotid artery, small arteries, venous lacunae, and nerve fibres.

The maxillary artery, with a measured wall thickness of ∼0.3–0.6 mm, ran ventral to the ear canal and gave rise to the mandibular alveolar artery, which entered the mandibular canal, as well as venous branches to the connective tissue of the ear and jaw attachment (Fig 10). The maxillary vein also contributed to a fibrovascular plexus that partially wrapped around the tympanic bulla, transitioned into the mandibular plexus within the IMFB, and formed a PAVR around the mandibular artery. On DICE-µCT, small branches of the maxillary vein were observed connecting to the intramandibular venous plexus.

**Figure 10.**
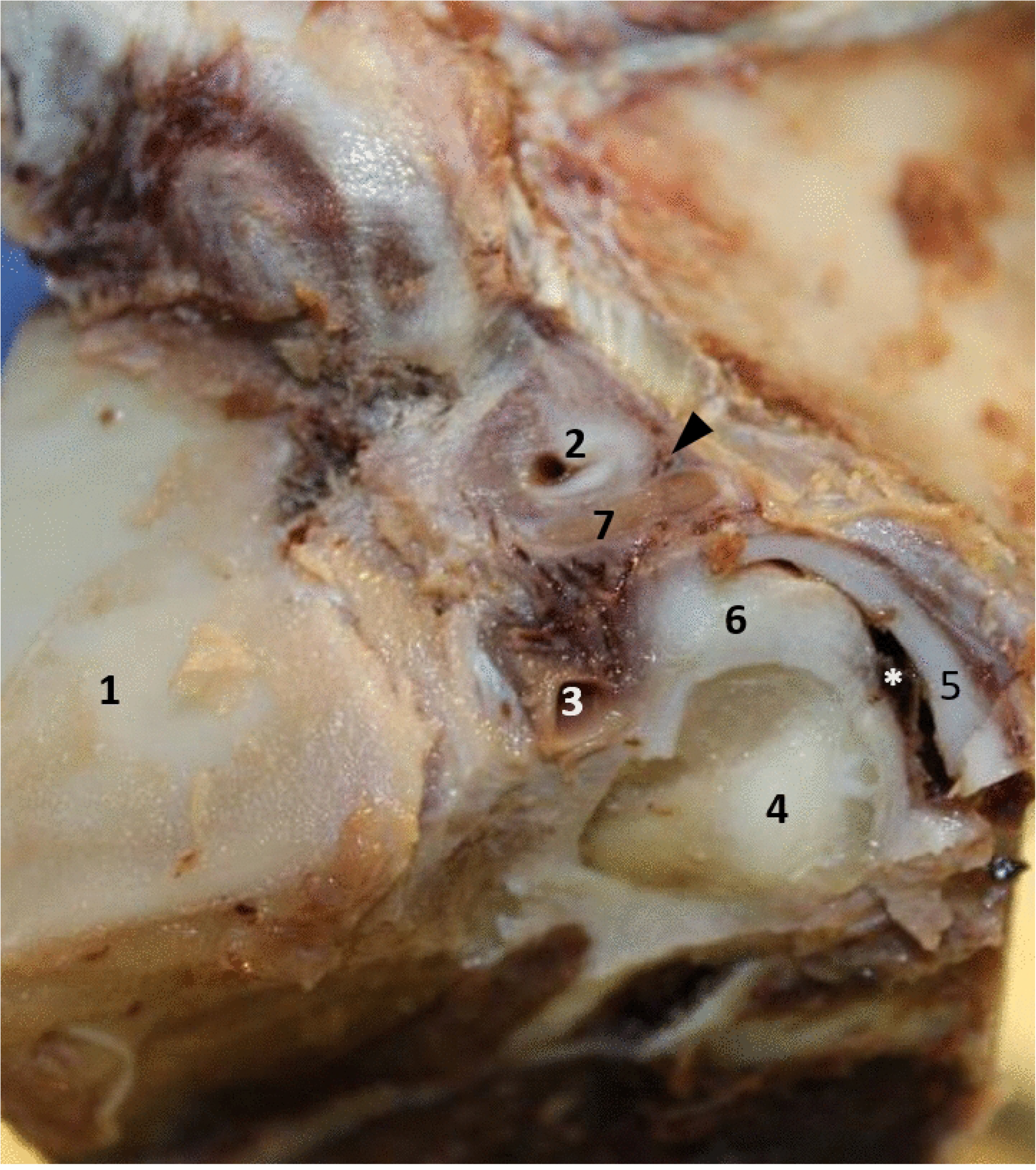
Left lateral view of a macroscopic dissection of the region of the external ear canal in striped dolphin, dissected to the level of the mandible. (1). 2. external ear canal; 3, Mandibular vein with the mandibular fibrovenous plexus dorsal to it, similar to the veins located caudal to the ear canal (arrowhead); 4, caudal most part of the acoustic fat body complex, close to the location of the tympanic bone; 5, cartilage at the medial end of the stylohyoid; 6, connective tissue surrounding the fat body channel; 7, facial nerve running ventral to the ear canal. The ventral scalenus muscle has been removed. Asterisk, posterolateral lobe of the peribullar sinus. The extramandibular fat body (EMFB) was partially removed but remains visible caudal to the mandible.

DICE-µCT showed the pterygoid, posterior, and middle sinus compartments, along with surrounding vascular structures. The venous plexuses formed continuous, extensive networks surrounding the sinuses and constituting a continuous interface with the tympanic bulla, periotic complex, and mandibular fat bodies. The distinction between sinus lumina and venous plexuses was not always possible in DICE-µCT, as their contours followed the same surfaces and displayed similar contrast, reflecting their highly complex morphology, specifically without blood pressure in a cadaver. The trabeculated air spaces and the relationships between sinuses, venous plexuses, and surrounding structures were intricately complex, and finer vascular and neural structures could not be traced confidently.

Histologically, the wall of the pterygoid sinus showed several small-to medium-sized nerve fibres (approximately 0.25 mm), as well as a ganglion within the loose connective tissue of the medial wall near the opening of the tuba auditiva. Numerous nerve fibres were observed macroscopically and microscopically in the ventrolateral wall of the pterygoid sinus, particularly at its border with the IMFB (Fig 11). The peribullar sinus wall contained several small-to medium-sized nerve fibres and many fine fibres. Sparse sensory corpuscles were present in the sinus walls, often associated with vascular structures. Similarly, the corpus cavernosum tympanicum showed dense innervation, forming a plexus of nerves, with scarce lamellar corpuscles and no other notable specialised nerve formations.

**Figure 11.**
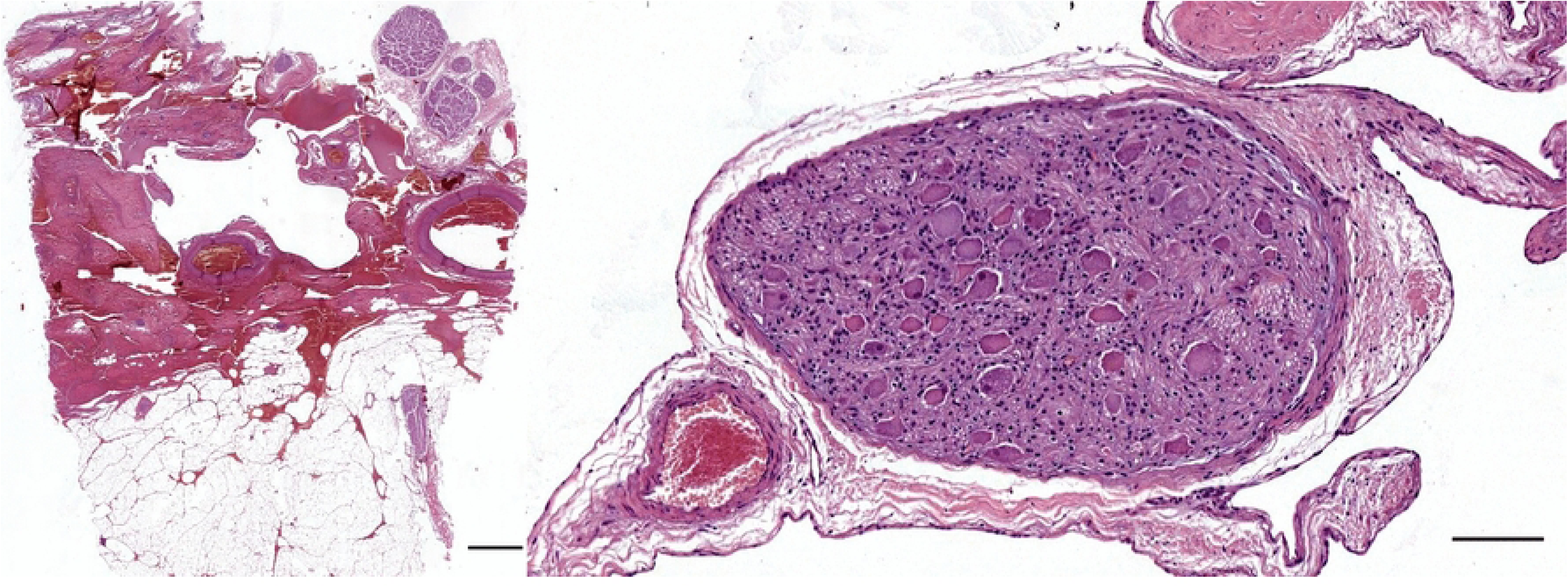
Left: HE-stained section through the ventrolateral wall of the pterygoid sinus, at the border with the IMFB (bottom) of a striped dolphin. Note the large nerve fibres in the fibrovenous wall. Scale bar 1 mm. Right: HE-stained section of a ganglion in the pterygoid sinus of a striped dolphin. Diameter: 0.7 x 0.4 mm. Scale bar 100 µm

## Discussion

The detailed morphology of the intra-and extramandibular fat bodies observed in this study reinforces and extends previous descriptions [27–31]. The findings highlight their continuous connection and close spatial relationships with both the external ear canal and middle ear. The posterolateral extension of the EMFB, positioned parallel to the external ear canal and flaring toward the blubber, provides morphological evidence for a lateral sound reception pathway complementary to the classical pan-bone window and gular routes [7,32]. The existence of a functionally significant lateral pathway remains debated. Cranford et al. [8] provided the strongest functional support through acoustic modelling in Ziphiidae, demonstrating that a lateral fat channel carries substantial acoustic energy to the tympanoperiotic complex. Yamato et al. [33] documented analogous lateral fat bodies in the minke whale, suggesting that such pathways vary considerably in geometry and functional role across lineages. However, caution should be applied before generalising lateral pathway models across odontocete taxa without species-specific acoustic and lipid biochemistry data. The posterolateral extension described here differs from the lateral fat channel of Ketten (1994) [32] in that it does not terminate at the pan bone but instead runs along the caudal mandibular margin, paralleling the ear canal before merging with the IMFB–EMFB complex and reaching the tympanoperiotic complex while bypassing the mandibular bone. Whether this extension constitutes a functionally significant acoustic pathway in delphinids, or whether it primarily serves a structural or anatomofunctional role associated with the external ear canal, cannot be determined from morphological evidence alone and requires acoustic modelling and species-comparative data. The microanatomical findings collectively suggest that the acoustic fat bodies are traversed by multiple neural elements, particularly branches of the mandibular and facial nerves, yet exhibit little evidence of terminal sensory specialisations, consistent with a sensitivity for local (such as vascular) regulation, supporting their interpretation as structurally integrated conduits for sound conduction rather than primary sensory or regulatory organs (cfr.[3]).

The cranial nerve pathways identified here confirm and expand upon previous observations of the V₃ and its branches [17,34]. The auriculotemporal nerve branched from the main trunk, together with mylohyoid and inferior alveolar nerve, and followed a trajectory caudal to the mandibular ramus, parallel to the external ear canal, with indications of a topographic connection to the facial nerve. Such anastomoses have been recently described in striped dolphin and dwarf sperm whale, although further rostral[35], while such anastomoses are known in other mammals[36,37]. The facial nerve’s topographical course, with small branches to the external ear canal musculature, is consistent with the description in delphinids[18,35]. The branches of V₃ (alveolaris inferior, monogastricus, pterygoid, mylohyoideus) followed expected courses including their relationships with surrounding fat and vascular structures, consistent with earlier reports (Rauschmann, 1992; Costidis & Rommel, 2012). Rare definitive sensory terminations within the acoustic fats or sinus walls were observed, suggesting these nerves act as conduits to more distant targets.

The masseter muscle was observed as a thin, reduced structure interspersed with the EMFB, consistent with its diminished functional role in odontocetes (Huggenberger, 2016). Its lack of a clear masseteric nerve in our material aligns with earlier descriptions of its poor development (Rauschmann, 1992), although Barthelmess (2017) reported a relatively large masseteric nerve in a Risso’s dolphin foetus macroscopic dissection.

Our findings confirm and refine previous descriptions of the venous plexuses (Boenninghaus, 1903; Fraser & Purves, 1960; Costidis & Rommel, 2012), documenting the full complement of intramandibular, peribullar, pterygoid, peritubal, mandibular alveolar, extramandibular, and corpus cavernosum plexuses. These venous networks were intimately associated with the sinuses, IMFB, and EMFB, with the IMFB showing the densest venous investment, with less dense vascularisation in the fat bodies’ core, supporting hypotheses of a specialised thermoregulatory or related physiological function (Costidis & Rommel, 2012). The corpus spongiosum tympanicum appeared as a dense venous structure embedded in the middle ear cavity, with abundant innervation but only sparse sensory specialisations, consistent with views of a venous compliance reservoir rather than an active erectile organ [3,38].

This study provides a detailed in-situ 3D reconstruction of the external ear canal’s downward spiral with associated fat, muscle and connective tissue, running ventral to the middle sinus, anterior to the posterior sinus (building on detailed descriptions of Purves [25]. At its medial end, the canal presented tissue of similar density to the acoustic fat bodies, although this should be interpreted with caution as tissue was contrast-enhanced for scanning. Facial nerve branches were observed at the ventral curvature of the canal, and the auriculotemporal nerve formed the main innervation pathway, consistent with its course in other delphinids [34].

Confocal microscopy revealed a dense and layered neurovascular architecture around the external ear canal, extending the sensory nerve formations’ tissue constitution [5], showing 3D insights on receptor morphology and providing a more comprehensive view of the ear canal’s elaborate sensory specialisations. Lamellar corpuscles were arranged longitudinally in the subepithelial layer and formed part of an interconnected circummeatal plexus, exhibiting diverse morphologies, including bifurcating, curving, or merging back into the nerve network. MCNs were consistently identified at the dermal–epidermal junction, with morphology and immunoprofiles (PGP9.5⁺/S100⁺/NF⁻) consistent with MCNs in terrestrial mammals [39–41]. The co-localisation of MCNs, presumed low-threshold, slowly adapting mechanoreceptors, with lamellar corpuscles, likely high-threshold, rapidly adapting sensors, although these may have evolved toward slower adaptation in the absence of a classical outer capsule [5], suggests a capacity for multimodal mechanosensation. This interpretation is further supported by the presence of intraepithelial PGP9.5⁺ structures lacking NF and S100 expression, whose morphology and distribution differ from known Langerhans cells and remain of uncertain identity. The identification of both deep and superficial neurovascular plexuses similar to the skin [42], and high-density innervation of arterioles and veins, reinforces the hypothesis that the canal serves not merely as a vestigial conduit, but as a specialised somatosensory interface. This structure may enable the dolphin to detect hydrodynamic stimuli, mechanical deformation, and local vascular changes, and potentially low-frequency acoustic stimuli, complementing the auditory roles of the tympano-periotic complex and acoustic fat bodies within a unified peripheral sensory system adapted to the aquatic environment.

Post-mortem collapse of venous spaces and filling by air in the sinuses, as observed in DICE-µCT, corroborates earlier suggestions that the air spaces expand into the venous spaces after death, making them difficult to distinguish [38,43].

Our findings corroborate and refine the described vascular architecture and functional interpretation, providing morphological indiations for the venous plexuses’ role as compliant reservoirs and extending the description of their neurovascular relationships.

## Conclusion

Collectively, this study demonstrates the anatomical and functional integration of acoustic, vascular, and neural structures in the dolphin auditory periphery, combining macroscopic dissection, DICE-µCT, histology, and confocal microscopy. The close association of fat bodies, venous plexuses, sinuses, and the external ear canal supports their role in sound reception, pressure regulation, and somatosensory feedback. Novel findings, including a complex neurovascular architecture and previously undescribed intraepithelial structures, suggest that the external ear canal may function as a multimodal sensory interface rather than a vestigial conduit. While limitations in resolving linking macro-and microanatomy, for tracing nervous and vascular structures, remain, these results refine existing anatomical models to further investigate the sensory physiology of cetaceans through classical methods, neurotracing, functional imaging, and biomechanical modelling.

## Acknowledgements

Part of this study was funded by the Italian Fondazione Cassa di Risparmio del Veneto e Rovigo, while another as part of the Project PCI2022-135022-2 financed by MICIU/AEI /10.13039/501100011033 and by the European Union NextGenerationEU/PRTR, supported by JPI Oceans. Gmar is funded by a Catalan Government Grups de Recerca Consolidats grant (ref. 2021 SGR 01195). The authors would like to acknowledge the Italian Zooprofilactic Institutes of Venezia, Lazio e Toscana, Lìguria, and Sardinia for collaboration in the sampling network, and Mariano Domingo (Department of Health and Anatomy, Faculty of Veterinary Medicine, Autonomous University of Barcelona, Bellaterra). Also thanks to Prof. dr. Hansjörg Schröder of the Department II of Anatomy (Neuroanatomy), University of Cologne (50924 Cologne, Germany) for allowing SDV to perform part of the macroscopic studies on site.

## Author contributions

Conceptualization: SDV, JMG, SM, MF, FL, CC, MA; Data Curation: SDV, MF, FL; Investigation: SDV, JMG, SH, MF, FL, CC, AF, MD, JF; Supervision: SM, MA; **Visualization**: SDV; Writing – Original Draft Preparation: SDV; Writing – Review & Editing: SDV, JMG, SM, SH, MF, FL, CC, AF, MCD, JF, MA

## References

1. Cranford T, Krysl P. Sound Paths, Cetaceans. Encyclopedia of Marine Mammals. Elsevier; 2018. pp. 901–904. doi:10.1016/B978-0-12-804327-1.00236-3

2. Ketten DR. Structure and Function in Whale Ears. Bioacoustics. 1997;8: 103–135.

3. Costidis A, Rommel SA. Vascularization of Air Sinuses and Fat Bodies in the Head of the Bottlenose Dolphin (*Tursiops truncatus*): Morphological Implications on Physiology. Frontiers in Physiology. 2012;3: 1–23. doi:10.3389/fphys.2012.00243

4. McCormick JG, Wever EG, Palin J, Ridgway SH. Sound Conduction in the Dolphin Ear. The Journal of the Acoustical Society of America. 1970;48: 1418–1428. doi:10.1121/1.1912302

5. De Vreese S, André M, Cozzi B, Centelleghe C, van der Schaar M, Mazzariol S. Morphological Evidence for the Sensitivity of the Ear Canal of Odontocetes as shown by Immunohistochemistry and Transmission Electron Microscopy. Sci Rep. 2020;10: 1–17. doi:10.1038/s41598-020-61170-4

6. Fraser FC, Purves PE. Hearing in cetaceans: Evolution of the accesory air sacs and the structure and funtion of the outer and middle ear in recent cetaceans. Bulletin of the British Museum (Natural History) Zoology. 1960;7: 1–140.

7. Norris KS. Evolution of Acoustic Mechanisms in Odontocetes. In: Drake ET, editor. Evolution and Environment. New Haven: Yale University Press; 1968. pp. 297–324.

8. Cranford TW, Krysl P, Hildebrand JA. Acoustic pathways revealed: simulated sound transmission and reception in Cuvier’s beaked whale (*Ziphius cavirostris*). Bioinspiration & Biomimetics. 2008;3: 016001. doi:10.1088/1748-3182/3/1/016001

9. Wei C, Erbe C, Smith AB, Yang W-C. Validated 3D finite-element model of the Risso’s dolphin (*Grampus griseus*) head anatomy demonstrates gular sound reception and channelling through the mandibular fats. Bioinspir Biomim. 2024;19: 056025. doi:10.1088/1748-3190/ad7344

10. Popov V, Supin A. Localization of the Acoustic Window at the Dolphin’s Head. Sensory Abilities of Cetaceans. Springer, Boston, MA; 1990. pp. 417–426. doi:10.1007/978-1-4899-0858-2_28

11. Goodson AD, Klinowska M. A Proposed Echolocation Receptor for the Bottlenose Dolphin (*Tursiops truncatus*): Modelling the Received Directivity from Tooth and Lower Jaw Geometry. In: Thomas JA, Kastelein RA, editors. Sensory Abilities of Cetaceans: Laboratory and Field Evidence. Boston, MA: Springer US; 1990. pp. 255–267. doi:10.1007/978-1-4899-0858-2_15

12. McCormick J. Anatomical, electrophysiological, and histological studies of dolphin auditory system: Establishment of a theory of hearing for dolphins. The Journal of the Acoustical Society of America. 2008;124: 2464. doi:10.1121/1.4782686

13. Romero-Vivas E, Leon-Lopez B. Analogy of the dolphin jaw to a metamaterial leaky wave antenna for sound directional detection. Journal of Sound and Vibration. 2025;604: 118987. doi:10.1016/j.jsv.2025.118987

14. Mooney TA, Nachtigall PE, Castellote M, Taylor KA, Pacini AF, Esteban J-A. Hearing pathways and directional sensitivity of the beluga whale, *Delphinapterus leucas*. Journal of Experimental Marine Biology and Ecology. 2008;362: 108–116. doi:10.1016/j.jembe.2008.06.004

15. Popov VV, Supin AYa, Klishin VO, Tarakanov MB, Pletenko MG. Evidence for double acoustic windows in the dolphin, *Tursiops truncatus*. The Journal of the Acoustical Society of America. 2008;123: 552–560. doi:10.1121/1.2816564

16. Fernández A, Edwards JF, Rodriguez F, De Los Monteros AE, Herraez P, Castro P, et al. “Gas and fat embolic syndrome” involving a mass stranding of beaked whales (family Ziphiidae) exposed to anthropogenic sonar signals. Veterinary Pathology. 2005;42: 446–457.

17. Rauschmann MA. Morphologie des Kopfes beim Schlanken Delphin *Stenella attenuata* mit besonderer Berücksichtigung der Hirnnerven. Makroskopische Präparation und moderne bildgebende Verfahren - Dissertation. Johann Wolfgang Goethe-Universität. 1992.

18. Cozzi B, Huggenberger S, Oelschläger HHA. Anatomy of Dolphins. Academic Press; 2017. Available: https://www.elsevier.com/books/anatomy-of-dolphins/cozzi/978-0-12-407229-9

19. IJsseldijk LL, Brownlow A, Mazzariol S. Best practice for cetacean post mortem investigation and tissue sampling. ASCOBANS/AC25/Inf32/Rev1. 2019; 71.

20. Cardona A, Saalfeld S, Schindelin J, Arganda-Carreras I, Preibisch S, Longair M, et al. TrakEM2 Software for Neural Circuit Reconstruction. PLOS ONE. 2012;7: e38011. doi:10.1371/journal.pone.0038011

21. Cardona A, Schindelin J, Eglinger J, Saalfeld S, Arganda-Carreras I. Register Virtual Stack Slices. In: ImageJ [Internet]. 2017 [cited 14 May 2020]. Available: https://imagej.net/Register_Virtual_Stack_Slices

22. Kikinis R, Pieper SD, Vosburgh KG. 3D Slicer: A Platform for Subject-Specific Image Analysis, Visualization, and Clinical Support. In: Jolesz FA, editor. Intraoperative Imaging and Image-Guided Therapy. New York, NY: Springer; 2014. pp. 277–289. doi:10.1007/978-1-4614-7657-3_19

23. Cignoni P, Callieri M, Corsini M, Dellepiane M, Ganovelli F, Ranzuglia G. MeshLab: an Open-Source Mesh Processing Tool. The Eurographics Association; 2008. doi:10.2312/LocalChapterEvents/ItalChap/ItalianChapConf2008/129-136

24. B.O. Community. Blender - a 3D modelling and rendering package. Amsterdam: Stichting Blender Foundation; 2018. Available: Retrieved from http://www.blender.org

25. Purves PE. Anatomy and Physiology of the Outer and Middle Ear in Cetaceans. In: Norris KS, editor. Whales, Dolphins and Porpoises. Berkeley, California, USA: University of California Press; 1966. pp. 320–380. Available: https://porpoise.org/library/anatomy-physiology-outer-middle-ear-cetaceans/

26. De Vreese S, André M, Mazzariol S. Morphology of the external ear canal in toothed whales. Proc Mtgs Acoust. 2019;37: 010016. doi:10.1121/2.0001281

27. Cranford TW, Amundin M, Norris KS. Functional morphology and homology in the odontocete nasal complex: implications for sound generation. J Morphol. 1996;228: 223–285. doi:10.1002/(SICI)1097-4687(199606)228:3<223::AID-JMOR1>3.0.CO;2-3

28. Harper CJ, McLellan WA, Rommel SA, Gay DM, Dillaman RM, Pabst DA. Morphology of the melon and its tendinous connections to the facial muscles in bottlenose dolphins (*Tursiops truncatus*). Journal of Morphology. 2008;269: 820–839. doi:10.1002/jmor.10628

29. Koopman HN, Iverson SJ, Read AJ. High concentrations of isovaleric acid in the fats of odontocetes: variation and patterns of accumulation in blubber vs. stability in the melon. Journal of Comparative Physiology B. 2003;173: 247–261.

30. Norris KS. The echolocation of marine mammals. The Biology of Marine Mammals. New York, NY: Academic Press; 1969. pp. 391–423.

31. Norris KS, Harvey GW. Sound transmission in the porpoise head. The Journal of the Acoustical Society of America. 1974;56: 659–664. doi:10.1121/1.1903305

32. Ketten DR. Functional Analyses of Whale Ears: Adaptations for Underwater Hearing. OCEANS’94’Oceans Engineering for Today’s Technology and Tomorrow’s Preservation’Proceedings. IEEE; 1994. pp. 264–270. Available: http://ieeexplore.ieee.org/abstract/document/363871/

33. Yamato M. The auditory system of the minke whale (*Balaenoptera acutorostrata*): a potential fatty sound reception pathway in a mysticete cetacean. Doctoral thesis, Massachusetts Institute of Technology and Woods Hole Oceanographic Institution. 2012. doi:10.1575/1912/5661

34. Barthelmess NG. Topographic Anatomy and Course of Cranial Nerves in the Risso’s Dolphin (*Grampus griseus*). Bachelor thesis, University of Cologne. 2017.

35. Nishimaniwa K, Yamada TK, Sekiya S, Amano M, Tajima Y. Morphological features of the orbicularis oculi muscle and facial nerve in four odontocete families, with comparisons within Cetartiodactyla. J Vet Med Sci. 2026;88: 1–12. doi:10.1292/jvms.25-0321

36. Schaller O, Schaller O. Illustrated Veterinary Anatomical Nomenclature. 4., aktualisierte Auflage. Constantinescu GM, editor. Stuttgart: Thieme; 2018.

37. Kadrie A, Toomey P, Callaway J, Gillespie MB, Boughter Jr. JD. The Auriculotemporal Nerve: A Comprehensive Review of Its Anatomical Variation and Clinical Manifestations. Laryngoscope Investigative Otolaryngology. 2025;10: e70238. doi:10.1002/lio2.70238

38. Boenninghaus G. Das Ohr des Zahnwales, zugleich ein beitrag zur theorie der Schalleitung: eine biologische studie. Zoologische Jahrbücher Abteilung für Anatomie und Ontogenie der Tiere. 1903;19: 189–360.

39. Dalsgaard C-J, Rydh M, Högerstrand A. Cutaneous innervation in man visualized with protein gene product 9.5 (PGP 9.5) antibodies. Histochemistry. 1989;92: 385–390. doi:10.1007/BF00492495

40. Ramírez GA, Rodríguez F, Quesada Ó, Herráez P, Fernández A, Espinosa-de-los-Monteros A. Anatomical Mapping and Density of Merkel Cells in Skin and Mucosae of the Dog: Location and Density of Canine Merkel Cells. The Anatomical Record. 2016;299: 1157–1164. doi:10.1002/ar.23387

41. Hoffman BU, Baba Y, Griffith TN, Mosharov EV, Woo S-H, Roybal DD, et al. Merkel Cells Activate Sensory Neural Pathways through Adrenergic Synapses. Neuron. 2018;100: 1401–1413.e6. doi:10.1016/j.neuron.2018.10.034

42. Eldridge SA, Mortazavi F, Rice FL, Ketten DR, Wiley DN, Lyman E, et al. Specializations of somatosensory innervation in the skin of humpback whales (*Megaptera novaeangliae*). The Anatomical Record. 2022;305: ar.24856. doi:10.1002/ar.24856

43. Costidis AM. The morphology of the venous system in the head and neck of the bottlenose dolphin (*Tursiops truncatus*) and Florida manatee (*Trichetus manatus latirostris*). PhD Thesis, University of Florida. 2012.

